# Functional connectivity within and between *n*-back modulated regions: An adult lifespan PPI investigation

**DOI:** 10.1101/2020.06.11.145219

**Authors:** Ekarin E. Pongpipat, Kristen M. Kennedy, Chris M. Foster, Maria A. Boylan, Karen M. Rodrigue

## Abstract

Working memory (WM) and its BOLD-related parametric modulation under load decrease with age. Functional connectivity (FC) generally increases with WM load; however, how aging impacts connectivity and whether this is load-dependent, region-dependent, or associated with cognitive performance is unclear. This study examines these questions in 170 healthy adults (*M*_*age*_ = 52.99 ± 19.18) who completed fMRI scanning during an *n*-back task (0-, 2-, 3-, and 4-back). FC was estimated utilizing a modified generalized psychophysiological interaction approach with seeds from fronto-parietal (FP) and default mode (DM) regions that modulated to *n*-back difficulty. FC analyses focused on both connectivity during WM engagement (task vs control) and connectivity in response to increased WM load (linear slope across conditions). Each analysis utilized within- and between-region FC, predicted by age (linear or quadratic), and its associations with in- and out-of-scanner task performance. Engaging in WM either generally (task vs control) or as a function of difficulty strengthened integration within- and between-FP and DM regions. Notably, these task-sensitive functional connections were robust to the effects of age. Stronger negative FC between FP and DM regions was also associated with better WM performance in an age-dependent manner, occurring selectively in middle- and older-adults. These results suggest that FC is critical for engaging in cognitively demanding tasks, and its lack of sensitivity to healthy aging may provide a means to maintain cognition across the adult lifespan. Thus, this study highlights the contribution of maintenance in brain function to support working memory processing with aging.

**Impact Statement:** The literature examining functional connectivity (FC) during working memory (WM) in healthy adults is mixed in both age effects and its relationship to performance. This study contributes to the literature by examining a large, adult lifespan sample, increased levels of WM load, and additional investigation of connections within and between fronto-parietal and default mode regions. Results revealed age-invariant strengthened FC during WM, suggesting that healthy aging may be resilient to FC changes. Additionally, negative FC between regions was associated with better WM performance in middle-aged and older adults, highlighting the important of FC maintenance to support successful WM ability.

## Introduction

Working memory (WM) ability, generally defined as the cognitive process that involves temporarily storing and manipulating information (Baddeley, 2000; Baddeley & Hitch, 1974), decreases with aging (Artuso, et al., 2017; Dobbs & Rule, 1989; Dumas, et al., 2001; Park et al., 2002). WM relies on a network of regions bilaterally in the frontal (e.g., lateral and medial prefrontal cortex and frontal operculum) and parietal lobes (e.g., intraparietal sulcus) (Owen, et al., 2005; Rottschy et al., 2012); however, the way in which these brain regions support WM across the adult lifespan is poorly understood. fMRI methods allow for the estimation of brain activation or connectivity under varying WM demand. Activation studies provide insight into the differential activation of brain regions during cognitive performance, whereas connectivity studies indicate how the brain synchronizes activity over time. Each method provides unique, but complementary, information about how the brain supports WM and maintains WM performance over time. Activation studies often report robust age differences in brain activation as WM demand increases, however, task-based connectivity studies are more mixed in the reporting of age effects, especially as related to supporting cognition.

Brain activation has been shown to modulate as a function of WM load increase (Owen et al., 2005; Kennedy et al, 2017; Rottschy et al., 2012; Wager & Smith, 2003). In addition, both positive modulation (i.e., increasing activity) of fronto-parietal (FP) regions and negative modulation (i.e., decreasing activity) of default mode (DM) regions to increasing WM load are reduced with increasing age (e.g., Kennedy et al., 2017), which is in accord with the Default-Executive Coupling Hypothesis of Aging (DECHA) model (Turner & Spreng, 2015). DECHA posits that coupling of activation in FPN regions alongside suppression of DMN regions during executive function tasks (including WM), is altered with aging, possibly resulting in poorer cognitive performance.

Maintaining synchronous activity over time (e.g., functional integration) is also critical to supporting complex cognitive functions. Functional integration of brain regions during a given cognitive task is necessary for proper task performance, and improper coupling of regions or networks likely underlies individual differences in performance (Shine et al., 2016). Functional connectivity (FC), a proxy for functional integration, has been explored during WM and demonstrates increases both within and between FP and DM regions under greater WM load (Di & Biswal, 2018; Hakun, et al., 2015; Heinzel et al., 2014; Heinzel et al., 2017; Nagel et al., 2011; Newton et al., 2011). The sensitivity of connectivity to task difficulty suggests that FC is an essential brain function that supports complex cognition (Smith, Gseir, Speer, & Delgado, 2016). Thus, it is critical to understand FC differences across the adult lifespan and further, how this brain property relates to cognitive performance.

Despite the consistent finding that connectivity in the FPN and DMN are responsive to task demands, age group comparison studies of younger vs. older adults have revealed differential patterns of FP connectivity to increasing WM load (Hakun, et al., 2015; Heinzel et al., 2014; Heinzel et al., 2017; Honey et al., 2002; Nagel et al., 2011). For instance, during the *n*-back paradigm across studies, older adults evidence decreased connectivity in FP regions during the 3-back condition (Heinzel et al., 2014, 2017), decreased FC for 3-back compared to 1-back conditions (Heinzel et al., 2014; Nagel et al., 2011), and also *increased* connectivity during 2-back condition (Heinzel et al., 2014). Similarly, the few studies that examine changes in functional connectivity to increasing task difficulty between FP-DM regions with regard to aging are equivocal, reporting both weakened negative FC (anti-phase synchronization) and reversed FC (strengthened in-phase synchronization) with aging (Hakun, Zhu, Johnson, & Gold, 2015; Turner & Spreng, 2015).

In addition to these FC findings, we recently reported that the significant inverse correlation between positive modulation clusters (which included largely FP regions) and negative modulation clusters (which included largely DM regions) to increased WM load was invariant to aging (Kennedy et al., 2017). We interpreted this functional coupling (i.e., strengthened positive modulation coupled with weakened negative modulation), as an individual difference measure that gauged the correlated modulation of these FP and DM regions, as would be predicted by the DECHA framework (Turner & Spreng, 2015). Importantly, that coupling was derived from a correlation of modulated activation and is not a direct measure of functional connectivity. However, the finding provides indirect support for an age-invariant property of network coupling in healthy aging (e.g., between FP and DM regions).

Finally, an important component of understanding the broader significance of aging effects on task-based connectivity is evidence for reliable association between FC and cognitive performance. To date, the literature on connectivity among fronto-parietal regions, age, and cognitive performance reports positive associations across the adult lifespan, positive associations in younger, but not older adults, as well as null findings (Heinzel et al., 2014; Heinzel et al, 2017; Nagel et al., 2011). Further, the impact of aging on *between* FP-DM coupling during WM with WM performance is unexamined. Thus, a gap exists in the understanding of this critical metric of brain function as it relates to aging and to cognitive performance.

Therefore, the present study examines functional connectivity (using modified generalized psychophysiological interactions; gPPI) both within- and between-FP and DM regions that previously demonstrated parametrically modulated activation in response to working memory load during an *n*-back task (in Kennedy et al., 2017). We computed PPI models for two contrasts: WM task compared to control conditions (2-3-4 vs 0-back), and increasing working memory load across conditions (2-3-4 back slope). For each of these contrasts, we sought to determine 1) the pattern of within- and between FP and DM region connectivity, 2) whether these within- and/or between-region connectivity patterns differ with age (linearly or non-linearly), and 3) whether within- and/or between-region connectivity was predictive of working memory performance (measured both inside and outside-of-scanner) across the adult lifespan. Based on the limited literature, we expected 1) strengthened positive connectivity within-FP and within-DM regions, (i.e., in-phase synchronization), and strengthened negative connectivity between FP and DM regions (i.e., anti-phase synchronization) when engaged in the task and as the task *n*-back load increased; and 2) given the limited and mixed associations of FC to aging and to cognitive performance, we suspected that FC could be age-invariant, but that its association to cognitive performance may be load- and/or age-dependent.

## Materials and Methods

### Participants

Participants consisted of 170 healthy adults aged 20 to 94 years (*M* = 52.99, *SD* = 19.18) recruited via media advertisements and flyers from the greater Dallas-Fort Worth metroplex. Exclusion criteria included cardiovascular disease, diabetes, cognition-altering medication, and a history of head trauma with loss of consciousness greater than 5 minutes, substance abuse, neurological, or psychiatric disorders. Participants were required to score > 25 on the Mini-Mental State Exam (MMSE; Folstein, Folstein, & McHugh, 1975) and ≤ 16 on the Center for Epidemiological Study Depression Scale (CES-D; Radloff, 1977). All participants were native English speakers, right-handed, had a minimum of high school education or equivalent, normal or corrected-to-normal vision, and provided informed consent according to the UT Southwestern Medical Center and UT Dallas institutional review boards. See Table 1 for participant demographics.

**Table 1.**
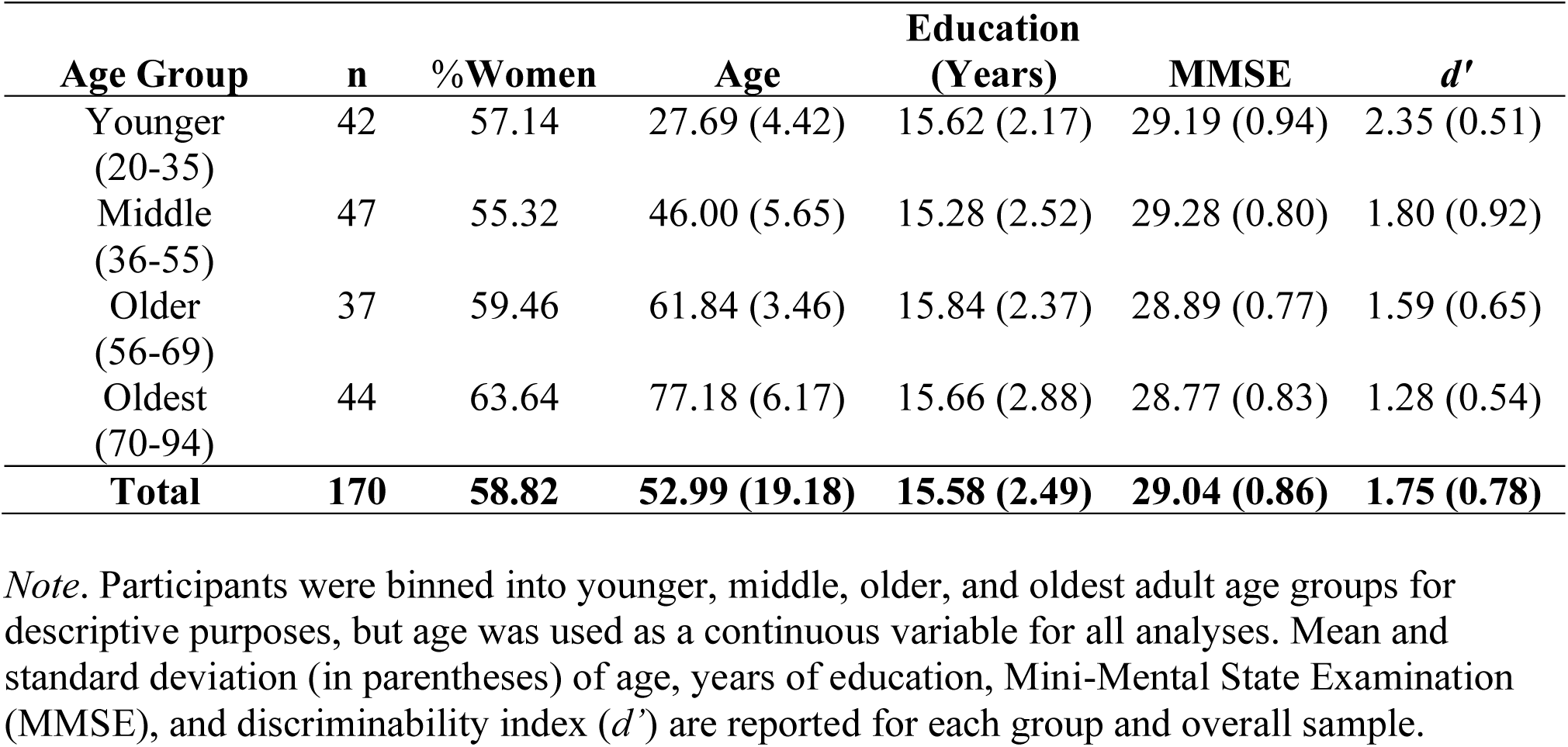
Participant Demographics by Age Group: Means (± Standard Deviation)

| **Age Group** | **n** | **% Women** | **Age** | **Education**<br>**(Years)** | **MMSE** | **$d'$** |
| --- | --- | --- | --- | --- | --- | --- |
| Younger<br>(20-35) | 42 | 57.14 | 27.69 (4.42) | 15.62 (2.17) | 29.19 (0.94) | 2.35 (0.51) |
| Middle<br>(36-55) | 47 | 55.32 | 46.00 (5.65) | 15.28 (2.52) | 29.28 (0.80) | 1.80 (0.92) |
| Older<br>(56-69) | 37 | 59.46 | 61.84 (3.46) | 15.84 (2.37) | 28.89 (0.77) | 1.59 (0.65) |
| Oldest<br>(70-94) | 44 | 63.64 | 77.18 (6.17) | 15.66 (2.88) | 28.77 (0.83) | 1.28 (0.54) |
| **Total** | **170** | **58.82** | **52.99 (19.18)** | **15.58 (2.49)** | **29.04 (0.86)** | **1.75 (0.78)** |

*Note.* Participants were binned into younger, middle, older, and oldest adult age groups for descriptive purposes, but age was used as a continuous variable for all analyses. Mean and standard deviation (in parentheses) of age, years of education, Mini-Mental State Examination (MMSE), and discriminability index ( $d'$ ) are reported for each group and overall sample.

### Procedures

Participants completed two cognitive assessment sessions followed by an MRI session, each completed on separate days and lasting approximately two hours, see Kennedy et al., (2017) for an unabridged description. During the cognitive sessions, participants completed a battery of cognitive tests that examined cognitive processes such as executive function, memory, problem-solving, and reasoning. During the MRI session, participants were trained on the in-scanner task followed by a collection of multimodal MRI images including functional and structural MR images.

### Working Memory (WAIS-DS)

During the cognitive sessions, participants completed the Digit Span subtest of the Wechsler Adult Intelligence Scale (WAIS-DS) (Wechsler, 2008). The WAIS-DS consisted of a forward, backward, and sequencing subsection where participants were given a series of numbers and were instructed to list the numbers in the same order as given (forward), in reverse order as given (backwards), or in numerical order (sequencing). Given that the digit span sequencing subtest was the most difficult task and most aligns with the *n*-back paradigm, this subtest was analyzed as a measure of out-of-scanner WM performance (Kennedy et al., 2017).

### MRI Acquisition

All participants were scanned on a single 3T Philips Achieva scanner equipped with a 32-channel head coil at the Advanced Imaging Research Center at the University of Texas Southwestern Medical Center. Functional data during an *n-*back task were collected using a T2*-weighted echo-planar imaging (EPI) sequence with 29 interleaved axial slices per volume providing full brain coverage and acquired parallel to the AC-PC line (64 × 64 × 29 matrix, 3.4 × 3.4 × 5 mm^3^, FOV = 220 mm^2^, TE = 30 ms, TR = 1.5 s, flip angle = 60°). High resolution structural images were also collected with a T1-weighted MP-RAGE sequence (160 sagittal slices, 1 × 1 x 1 mm^3^ voxel size; 256 × 204 x 160 matrix, TR = 8.3 ms, TE = 3.8 ms, flip angle = 12°).

### *N*-back Training

To ensure understanding of the *n-*back paradigm, participants were trained on the task prior to entering the scanner. Participants were first given instructions regarding the 0-back condition and were taught how to respond if a number was the same or different than an instructed number using an identical button box to that used in-scanner. Participants practiced the 0-back until they achieved greater than 80% accuracy. Participants were then trained similarly for the 2, 3, and 4-back conditions. After practicing each WM load condition, participants completed an abbreviated version of the full task.

### FMRI Task: Digit *n*-back

In the scanner, participants completed three functional runs of the *n*-back task in a block design. The task was presented and behavior recorded using PsychoPy software v1.77.02 (Peirce, 2007, 2009). Participants viewed the stimuli on a monitor mounted to the rear of the scanner, which was visible via a mirror mounted on the head coil. For each run, two blocks of each condition were presented in a pseudo-counterbalanced order. Each 0-back block included 10 trials, while each 2, 3, and 4-back block included 20 trials. For each block, a 5-sec cue indicated which *n*-back load was to follow (i.e., 0-, 2-, 3-, or 4-back). The cue was followed by a 2-sec fixation cross prior to the presentation of the digit. Digits 2-9 were pseudo-randomly presented for 500-ms with a 2000-ms inter-stimulus interval. Of the 420 trials, 144 (34.3%) were match (same) trials (i.e., 18 [4.2%] for 0-back and 42 [10.0%] each for 2, 3, and 4-back) and 276 (65.7%) were non-match (different) trials (i.e., 42 [10.0%] for 0-back and 78 [18.6%] each for 2, 3, and 4-back).

### FMRI Pre-Processing

Prior to pre-processing, each scan was visually inspected for quality and motion artifacts. A standard pre-processing pipeline using SPM8 software (Wellcome Department of Cognitive Neurology, London, UK) via Matlab R2012b (Mathworks) consisted of realignment, co-registration of functional to T1 anatomical images, warping functional images to MNI space using the T1 anatomical to MNI warp, and spatial smoothing of the functional images using an 8mm FWHM gaussian kernel. Artifact Repair Toolbox was also utilized to examine movement and intensity shift in the functional images (Mazaika, Whitfield, & Cooper, 2005). Volumes were marked as outliers if movement was greater than 2 mm of translation or 2 degrees of rotation, or an intensity shift greater than 3% deviation from the mean global intensity shift. Functional runs were excluded if 40 or more volumes (i.e., 15% of total volumes) were marked as outliers for movement. Only participants with at least two functional runs were included in the subsequent psychophysiological interaction (PPI) and group analyses. Six participants were excluded from further analyses: more than one functional run marked as an outlier (*n* = 3), poor T1-weighted scan acquisition (*n* = 2), provided no response to greater than 15% of the trials (*n* = 1).

### Regions of Interest (ROIs)

Seed regions of interest (ROIs) were chosen from the statistical map of the parametric modulation effects reported in Kennedy, et al. (2017), however, we limited the seeds to brain regions located primarily within fronto-parietal and default mode networks as these regions are consistently reported as being task sensitive in the working memory literature (see Table 2 and Figure 1). Fronto-parietal regions were chosen from peak global and local maxima from clusters showing positive modulation that lie within regions associated with FPN. Default mode regions were chosen from peak global and local maxima from clusters demonstrating negative modulation that are typically associated with DMN. Spheres with 6mm radius were created from each maxima coordinate. The seed ROIs were also used as target ROIs. Although these ROIs were extracted from the parametric modulation difficulty contrast, most of these regions were also found to be age-sensitive (exceptions included ROIs 9-12 within the FP ROIs). Confirmation that the selected seeds fell within the respective FP or DM network, a mask was created combining atlases from Neurosynth (Yarkoni, Poldrack, Nichols, Essen, & Wager, 2011), Yeo’s 7 Networks (Yeo et al., 2011), CAREN (Doucet, Lee, & Frangou, 2019), and Shirer’s 90 functional ROIs (Shirer, Ryali, Rykhlevskaia, Menon, & Greicius, 2012). For the Neurosynth atlas, the FPN mask combined a mask of the phrases: “working memory”, fronto parietal”, “frontoparietal”, and “frontoparietal network”, while the DMN mask combined a mask of the phrases: “default”, “default mode”, “default network”, “dmn”, and “network dmn”. All selected seeds fell within their respective networks, with the exception of ROI 15 (supramarginal gyrus) that fell outside the DMN mask; however, the pattern of results was identical with the inclusion or exclusion of this ROI.

**Table 2.** Regions of Interest: 6mm-radius Spheres used as both Seeds and Targets

| ROI | MNI |  |  | Network | Hem. | Region |
| --- | --- | --- | --- | --- | --- | --- |
|  | *x* | *y* | *z* |  |  |  |
| 1 | 42 | -42 | 42 | fronto-parietal | R | supramarginal gyrus |
| 2 | -39 | -48 | 42 | fronto-parietal | L | inferior parietal lobule |
| 3 | -30 | -60 | 42 | fronto-parietal | L | inferior parietal lobule |
| 4 | 36 | -54 | 42 | fronto-parietal | R | inferior parietal lobule |
| 5 | 33 | 12 | 54 | fronto-parietal | R | middle frontal gyrus |
| 6 | -27 | 6 | 54 | fronto-parietal | L | middle frontal gyrus |
| 7 | 45 | 33 | 24 | fronto-parietal | R | inferior frontal triangularis |
| 8 | -42 | 27 | 24 | fronto-parietal | L | inferior frontal triangularis |
| 9 | -39 | 51 | 6 | fronto-parietal | L | middle frontal gyrus |
| 10 | 36 | 60 | 6 | fronto-parietal | R | middle frontal gyrus |
| 11 | 39 | 30 | 48 | fronto-parietal | R | middle frontal gyrus |
| 12 | -36 | 27 | 48 | fronto-parietal | L | middle frontal gyrus |
| 13 | -3 | 60 | 3 | default | L | medial superior frontal gyrus |
| 14 | -6 | -51 | 27 | default | L | posterior cingulum |
| 15 | 48 | -30 | 24 | default | R | supramarginal gyrus |
| 16 | -6 | -15 | 42 | default | L | middle cingulum |
| 17 | 9 | -21 | 45 | default | R | middle cingulum |
| 18 | -12 | -27 | 42 | default | L | middle cingulum |
| 19 | 48 | 3 | -33 | default | R | inferior temporal gyrus |
| 20 | -45 | 0 | -36 | default | L | inferior temporal gyrus |

Note: These regions were chosen from peak global and local maxima from prior results of positive and negative parametric modulation of the *n*-back task within fronto-parietal and default mode regions (Kennedy et al., 2017). X, Y, and Z coordinates are in Montreal Neurological Institute (MNI) atlas space. Abbreviations: hemisphere (hem.), left (L), and right (R).

**Figure 1.**
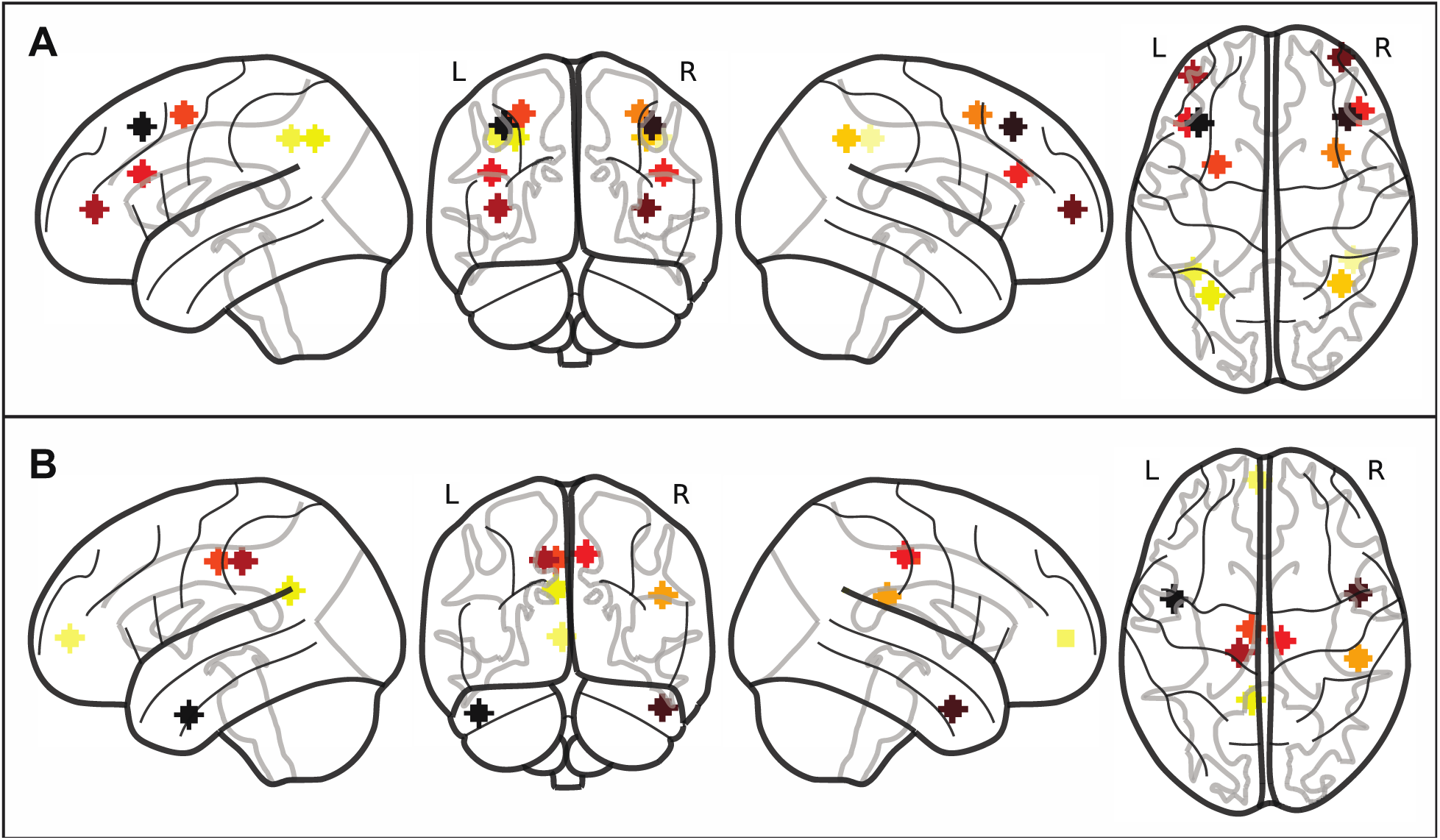
Sphere regions of interest within the fronto-parietal (A) and default mode (B) regions. Coordinates for the ROIs were chosen using the statistical map from the parametric modulation effects reported in Kennedy et al. (2017) using peak global and local maxima from regions that positively modulated within FP regions (A) and negatively modulated within DM regions (B) in response to working memory. Abbreviations: default-mode (DM), frontal-parietal (FP), inferior parietal lobule (IPL), inferior temporal gyrus (ITG), left (L), middle cingulate cortex (MCC), middle frontal gyrus (MFG), medial superior frontal gyrus (mSFG), posterior cingulate cortex (PCC), region of interest (ROI), right (R), supramarginal gyrus (SMG), and inferior frontal triangularis (TrIFG).

### Psychophysiological Interaction (PPI) Analyses

To examine functional connectivity, modified generalized psychophysiological interaction (gPPI) methods (Cisler, Bush, & Steele, 2014; Di, Reynolds, & Biswal, 2017; Di, Zhang, & Biswal, 2018; K J Friston et al., 1997; McLaren, Ries, Xu, & Johnson, 2012) were implemented in Analysis of Functional NeuroImages (AFNI) software (Chen, 2015; Cox, 1996) utilizing *a priori* orthogonal contrasts (Kaufman & Sweet, 1974; Lewis & Mouw, 1972). Second-level analyses were performed using the *stats* package and visualized using the *ggplot2* package in R via RStudio (R Core Team, 2019; RStudio Team, 2018; Wickham, 2016).

We utilized gPPI methods to examine whether the BOLD activity between two ROIs have a strengthened correlation (strengthened positive or negative correlation of activity across time) during a given task contrast using the pipeline illustrated in Figure 2. To create the psychological condition regressor, *n*-back working memory load (0-, 2-, 3-, 4-back), along with fixation, was treated as a categorical variable and *a priori* orthogonal contrasts codes were created. Specifically, contrasts were specified to test the effects of 1) *n*-back [0-, 2-, 3-, 4-back] compared to fixation [fixation: −0.800; 0-back: 0.200; 2-back: 0.200; 3-back: 0.200; 4-back: 0.200], 2) the task [2-, 3-, and 4-back] compared to control [0-back] condition [fixation: 0.000; 0-back: −0.750; 2-back: 0.250; 3-back: 0.250; 4-back: 0.250], 3) the linear parametric (slope) effect of task [fixation: 0.000; 0-back: 0.000; 2-back: −0.500; 3-back: 0.000; 4-back: 0.500, and 4) the quadratic parametric (slope) effect of task [fixation: 0.000; 0-back: 0.000; 2-back: −0.333; 3-back: 0.667; 4-back: −0.333]. Each psychological contrast was convolved using the canonical hemodynamic response function (HRF) from SPM. The convolved orthogonal contrast was used as the psychological variable (*PSY*). The mean BOLD signal was extracted from the seed ROI, the temporal trend (i.e., constant, linear, and quadratic) was removed from the time series, and then deconvolved using the same canonical HRF. The temporally detrended time series of the seed ROI was used as the physiological regressor (*PHYS*). To create each PPI regressor, the deconvolved physiological time series and each unconvolved orthogonal contrast were multiplied. The PPI term was then convolved using the same canonical HRF and used as the PPI variable (*PPI*). The *PSY, PHYS*, and *PPI* regressors were then concatenated across the functional runs. Each voxel’s concatenated BOLD time series was then regressed on the *PSY, PHYS*, and *PPI* variables while simultaneously controlling for temporal drift (i.e., baseline, linear, and quadratic), 24-motion parameters (Friston, Williams, Howard, Frackowiak, & Turner, 1996), the mean time series extracted from subject specific white matter (WM) and cerebral spinal fluid (CSF) masks, along with a 210s high-pass (HP) filter within a general linear model (GLM) framework (see Equation 1). The mean regression coefficient (i.e., unstandardized slope) for each *PPI* variable was extracted for each seed-to-target ROI (4 regression coefficients [*PPI*] * 20 seed ROIs * 19 target ROIs * = 1,520 regression coefficients per subject).

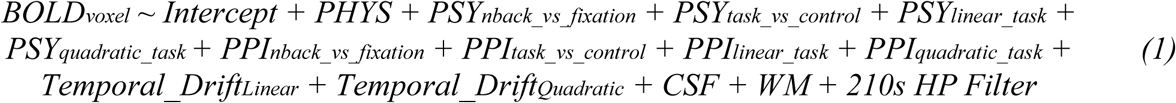

**Figure 2.**
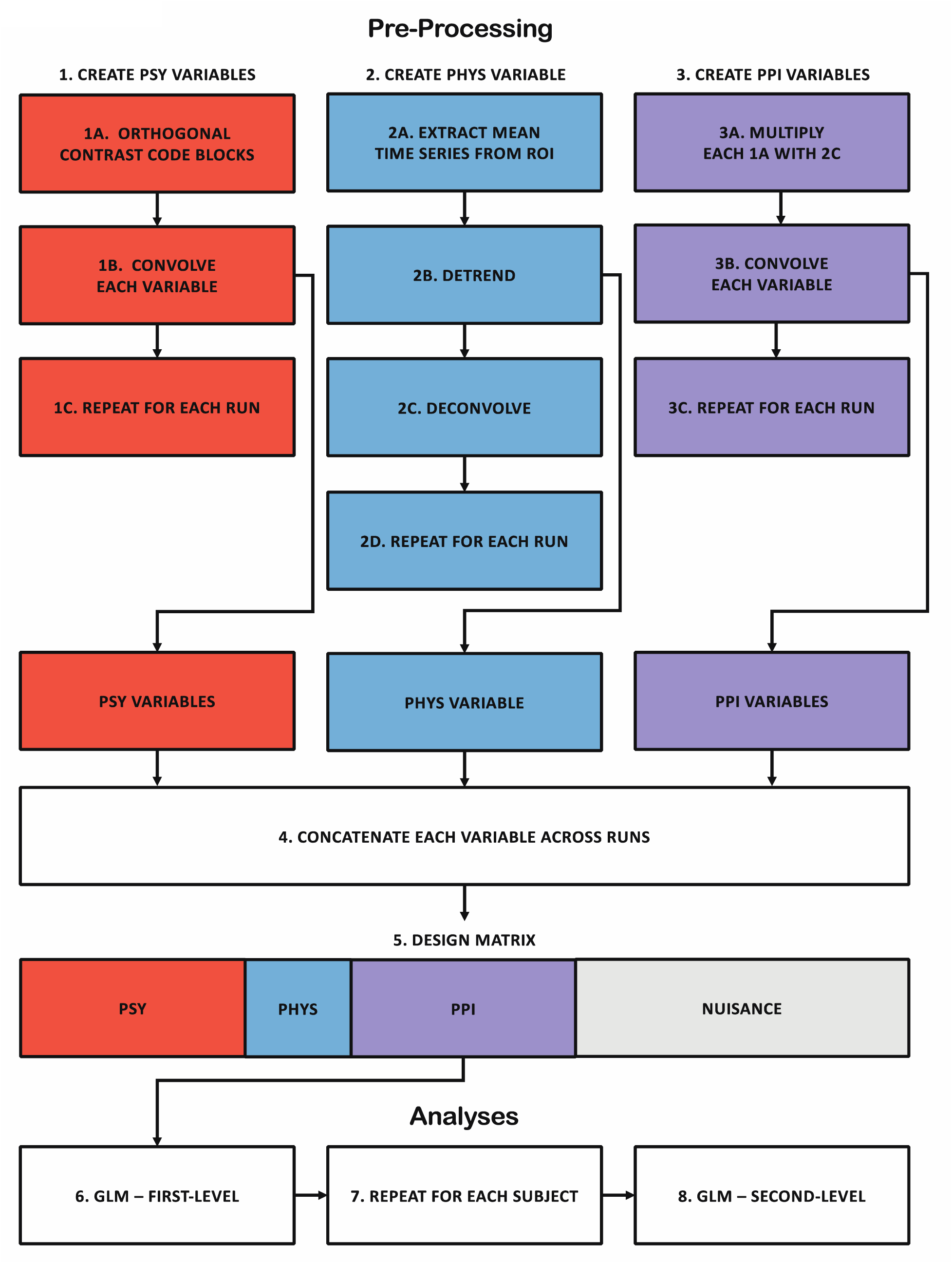
Psychophysiological Interaction (PPI) Analysis Pipeline. To create the *PSY* variable (Step 1), the time-series of the task was orthogonally-contrast coded (1A), each contrast code was then convolved (1B), and these steps were repeated for each run (1C). To create the *PHYS* variable (Step 2), the mean time-series was extracted from a ROI (2A), detrended linearly and quadratically (2B), deconvolved (2C), and these steps were repeated for each run (2C). To create the *PPI* variable (Step 3), each contrast code (1A) and deconvolved time-series (2C) was multiplied (3A), convolved (3B), and these steps were repeated for each run (3C). The *PSY* (1B), *PHYS* (2B), and *PPI* variable (3B) were concatenated across runs (4) and used as predictors along with nuisance variables (5) in the first-level GLM analysis (6). Nuisance variables in the GLM analysis included temporal drift, 24-motion parameters, and CSF and WM time-series along with a high-pass filter. These steps were then repeated for each subject (7). The estimates from the first-level analyses were then used in second-level analyses (8). This pipeline was repeated for each ROI. Abbreviations: general linear model (GLM), physiological (*PHYS*), psychological (*PSY*), and psychophysiological interaction (*PPI*).

### Group Analyses

To test FC within- and between the fronto-parietal and default mode regions, a group-level analysis was performed on each mean *PPI* regression coefficient of each seed-to-target ROI combination within a GLM framework (see Equation 2) with the exclusion of when the seed and target ROIs were identical. In other words, the 1,520 estimates were analyzed in an intercept-only model. Given the large number of non-independent analyses, the *α*-level (Type I error) was adjusted using the *Meff* correction (Derringer, 2018). The *Meff* value of all the *PPI* variables was calculated to be 1501.79. The *Meff* value was then used to divide by the overall *α* of 0.05 to obtain the corrected *α* of 0.00003.

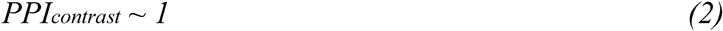

For subsequent analyses, *PPI* regression coefficients of each seed-to-target ROI were averaged to create FC within FP “network” (FPN), FC within DM “network” (DMN), and FC between FPN and DMN for each *PPI* variable (4 regression coefficients [*PPI*] * 3 FC pairs = 7 estimates per subject). The *Meff* correction for these analyses equaled 11.84. Using an overall *α* of 0.05, the corrected *α* used was 0.004. To test whether FC within and between the FPN and DMN varied across the adult lifespan, each *PPI* variable and FC pair was regressed on linear and quadratic age (see Equation 3). Linear age was mean-centered and quadratic age was the square of mean-centered age. To test whether there was an association between FC and *n*-back task performance across the adult lifespan, *n*-back performance using *d’* was regressed on FC of each *PPI* variable and network pair, age, and quadratic age along with its interactions (see Equation 4). FC of each *PPI* variable and network pair was mean-centered across participants. *d’* was calculated as z(hit rate) – z(false alarm rate). False alarm rates of 0 were adjusted using 1/(2N) where N was the total number of possible false alarms (i.e., *N* = 276), while hit rates of 1 were adjusted using 1-1/(2N) where N represented the total number of possible hits (i.e., *N* = 144). Lastly, to determine if the relationship between FC and WM performance was generalizable to out-of-scanner WM performance, digit span sequencing performance was regressed on FC of each *PPI* variable and network pair, age, and quadratic age along with its interactions (see Equation 5).

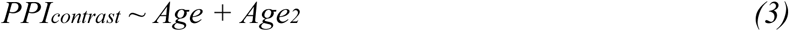

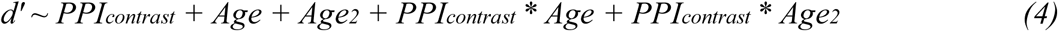

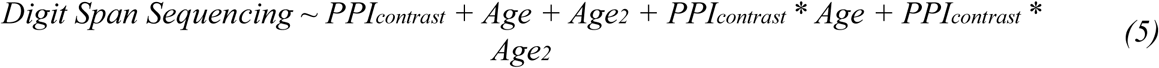

## Results

### Functional connectivity during *n*-back

To test for significant FC within- and between-FP and DM regions during *n*-back, the regression coefficient of each *PPI* variable and seed-to-target ROI combination was analyzed in an intercept-only model. Figure 3 displays both the unthresholded and thresholded results. Significant, thresholded results are summarized below as a range between the minimum regression coefficient (*b*) and its respective statistics (i.e., *t*-statistic and adjusted partial *r*^*2*^) to the maximum *b* and its respective statistics for each *PPI* variable and FC region.

**Figure 3.**
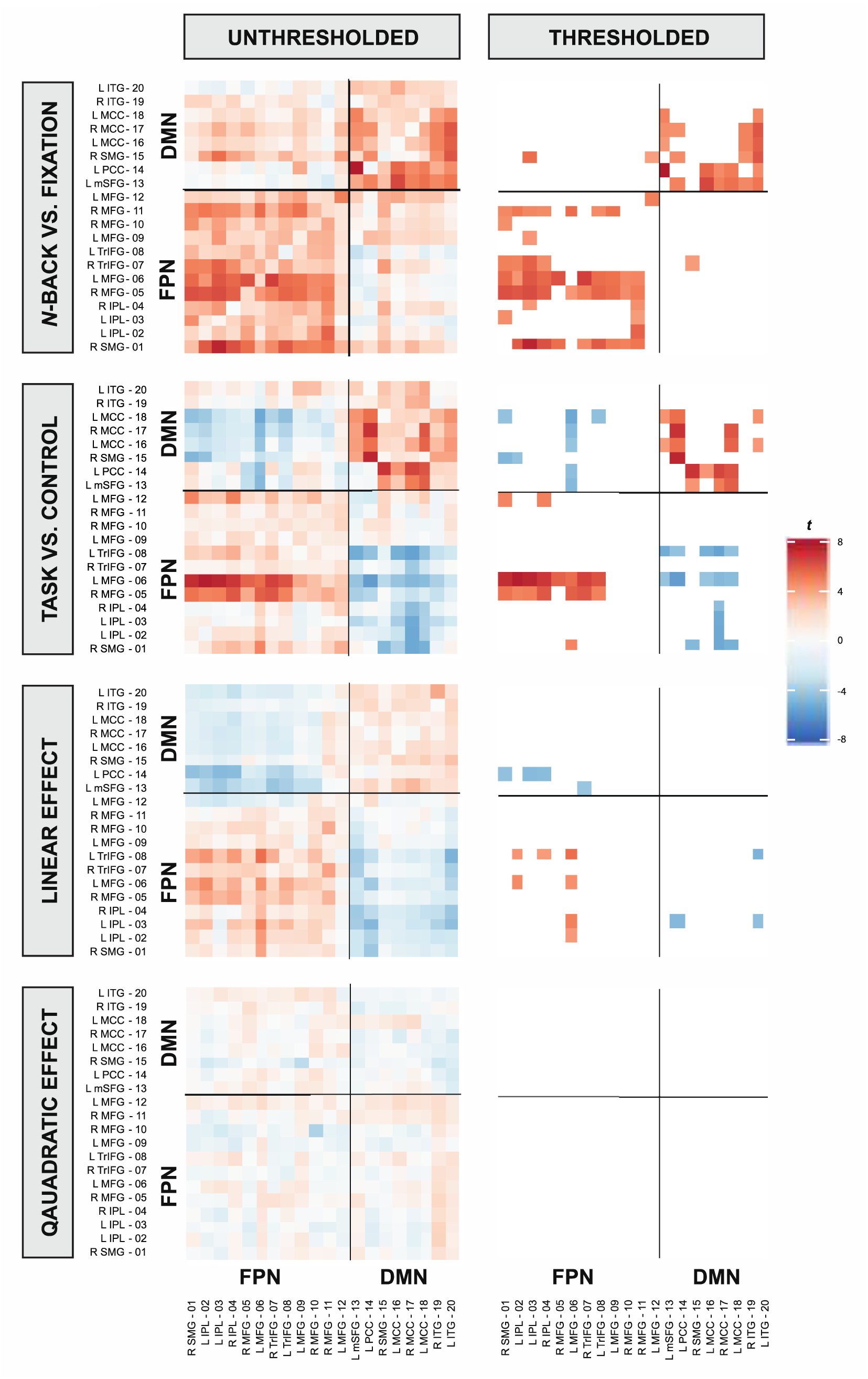
Unthresholded (A-D) and thresholded (E-H) t-statistics of each ROI pair for each PPI variable. Across *PPI* variables a generally negative correlation pattern is observed between FPN and DMN while a positive correlation pattern is displayed within-FPN and within-DMN (with the exception of the *n*-back vs fixation contrast). These effects were significant for the *n*-back vs. fixation, the task vs. control, and the linear effect of task contrasts. However, the positive correlation within-DM was not significant for the linear effect of task contrast. Significant effects were thresholded using an *α*_*Meff*_ of 0.00003. Abbreviations: default-mode network (DMN), fronto-parietal network (FPN), psychophysiological interaction (*PPI*), and region of interest (ROI). A full list of ROIs and their respective abbreviations can be seen in Table 1.

Regions within the FPN and within the DMN, respectively, showed significant positive FC (positive correlation; in-phase synchronization) during the *n*-back compared to fixation, (significant FC within-FPN: *b* = 0.20, *t*(169) = 5.18, adjusted *rp*^*2*^ = 0.14 to *b* = 1.55, *t*(169) = 5.68, adjusted *rp*^*2*^ = 0.16); significant FC within-DMN: *b* = 0.41, *t*(169) = 5.11, adjusted *rp*^*2*^ = 0.13 to *b* = 1.67, *t*(169) = 6.67, adjusted *rp*^*2*^ = 0.21), as well as during task compared to the control condition (significant FC within-FPN: *b* = 0.68, *t*(169) = 5.67, adjusted *rp*^*2*^ = 0.16 to *b* = 1.32, *t*(169) = 6.01, adjusted *rp*^*2*^ = 0.18; significant FC within-DMN: *b* = 0.48, *t*(169) = 4.57, adjusted *rp*^*2*^ = 0.11 to *b* = 2.86, *t*(169) = 7.24, adjusted *rp*^*2*^ = 0.24). Regions within FPN revealed significant strengthened positive FC as *n*-back load increased, but not within DMN (significant FC within-FP: *b* = 0.31, *t*(169) = 4.69, adjusted *rp*^*2*^ = 0.12 to *b* = 0.59, *t*(169) = 5.14, adjusted *rp*^*2*^ = 0.14). There were no significant quadratic task effects of FC within-FP or within-DM regions.

Regions between FP and DM networks revealed significant strengthened negative FC (negative correlation; anti-phase synchronization) during the *n*-back compared to fixation (significant: *b* = 0.57, *t*(169) = 4.28, adjusted *rp*^*2*^ = 0.10 to *b* = 0.98, *t*(169) = 4.36, adjusted *rp*^*2*^ = 0.10), during the task compared to the control condition (significant: *b* = −1.84, *t*(169) = −4.85, adjusted *rp*^*2*^ = 0.12 to *b* = −0.55, *t*(169) = −4.40, adjusted *rp*^*2*^ = 0.10), and during the linear effect of task (significant: *b* = −0.69, *t*(169) = −4.51, adjusted *rp*^*2*^ = 0.11 to b = −0.37, *t*(169) = −4.58, adjusted *rp*^*2*^ = 0.11). FC between FP and DM regions was not significant in the quadratic effect of task. Thus, regions within the FP and DM networks, along with regions between FP and DM networks were functionally connected to a greater extent during the task as compared to fixation or control, and additionally as a function of increasing *n*-back load.

### FC during *n*-back by Age

To determine whether FC within- and between-FPN and DMN during the *n*-back was moderated by age, each *PPI* variable and network pair (i.e., mean regression coefficient within-FPN, within-DMN, and between FPN-DMN) were regressed on linear and quadratic age (see Figure 4 and Table 3). Using a corrected *α* of 0.004, FC within- and between-FPN and DMN did not significantly change with linear (*p’s* > .083) or quadratic age (*p’s* > .020), suggesting that both within- and between-FP and DM connectivity during task were age-invariant.

**Table 3.** Second-Level Analysis of Each PPI Variable and FC by Linear and Quadratic Age

| **PPI Variable** | **FC** | **Variable** | ***b*** | ***SE*** | ***t*(167)** | ***p*** |  |
| --- | --- | --- | --- | --- | --- | --- | --- |
| Back vs.<br>Fixation | Within FPN | Intercept | 0.61 | 0.09 | 6.90 | < 0.001 | * |
|  |  | Age | 0.00 | 0.00 | -1.05 | 0.295 |  |
|  |  | Age2 | 0.00 | 0.00 | -0.56 | 0.573 |  |
|  | Within DMN | Intercept | 0.61 | 0.10 | 6.22 | < 0.001 | * |
|  |  | Age | 0.00 | 0.00 | -1.21 | 0.228 |  |
|  |  | Age2 | 0.00 | 0.00 | -0.13 | 0.896 |  |
|  | Between FPN and<br>DMN | Intercept | 0.26 | 0.11 | 2.50 | 0.013 |  |
|  |  | Age | 0.01 | 0.00 | 1.63 | 0.105 |  |
|  |  | Age2 | 0.00 | 0.00 | -0.62 | 0.535 |  |
| Task vs.<br>Control | Within FPN | Intercept | 0.32 | 0.11 | 3.05 | 0.002 | * |
|  |  | Age | 0.00 | 0.00 | -0.83 | 0.407 |  |
|  |  | Age2 | 0.00 | 0.00 | 0.27 | 0.786 |  |
|  | Within DMN | Intercept | 0.58 | 0.12 | 4.87 | < 0.001 | * |
|  |  | Age | 0.00 | 0.00 | -0.87 | 0.385 |  |
|  |  | Age2 | 0.00 | 0.00 | 0.51 | 0.613 |  |
|  | Between FPN and<br>DMN | Intercept | -0.21 | 0.11 | -1.82 | 0.071 |  |
|  |  | Age | 0.01 | 0.00 | 1.55 | 0.122 |  |
|  |  | Age2 | 0.00 | 0.00 | -0.98 | 0.330 |  |
| Linear Task | Within FPN | Intercept | 0.21 | 0.05 | 4.41 | < 0.001 | * |
|  |  | Age | 0.00 | 0.00 | 0.65 | 0.514 |  |
|  |  | Age2 | 0.00 | 0.00 | -2.36 | 0.020 |  |
|  | Within DMN | Intercept | 0.14 | 0.05 | 2.56 | 0.012 |  |
|  |  | Age | 0.00 | 0.00 | -2.30 | 0.022 |  |
|  |  | Age2 | 0.00 | 0.00 | -0.08 | 0.936 |  |
|  | Between FPN and<br>DMN | Intercept | -0.20 | 0.06 | -3.39 | 0.087 | * |
|  |  | Age | 0.00 | 0.00 | 1.41 | 0.160 |  |
|  |  | Age2 | 0.00 | 0.00 | 1.06 | 0.290 |  |
| Quadratic<br>Task | Within FPN | Intercept | 0.08 | 0.08 | 1.09 | 0.276 |  |
|  |  | Age | 0.00 | 0.00 | 1.64 | 0.103 |  |
|  |  | Age2 | 0.00 | 0.00 | -1.53 | 0.127 |  |
|  | Within DMN | Intercept | -0.04 | 0.08 | -0.42 | 0.674 |  |
|  |  | Age | 0.01 | 0.00 | 1.75 | 0.082 |  |
|  |  | Age2 | 0.00 | 0.00 | -0.58 | 0.566 |  |
|  | Between FPN and<br>DMN | Intercept | -0.01 | 0.09 | -0.12 | 0.904 |  |
|  |  | Age | 0.00 | 0.00 | 1.39 | 0.165 |  |
|  |  | Age2 | 0.00 | 0.00 | 0.87 | 0.387 |  |

*Note.* \*Significant effects; $p < \text{Meff-corrected } \alpha$ of 0.004. Abbreviations: default-mode network (DMN) and fronto-parietal network (FPN)

**Figure 4.**
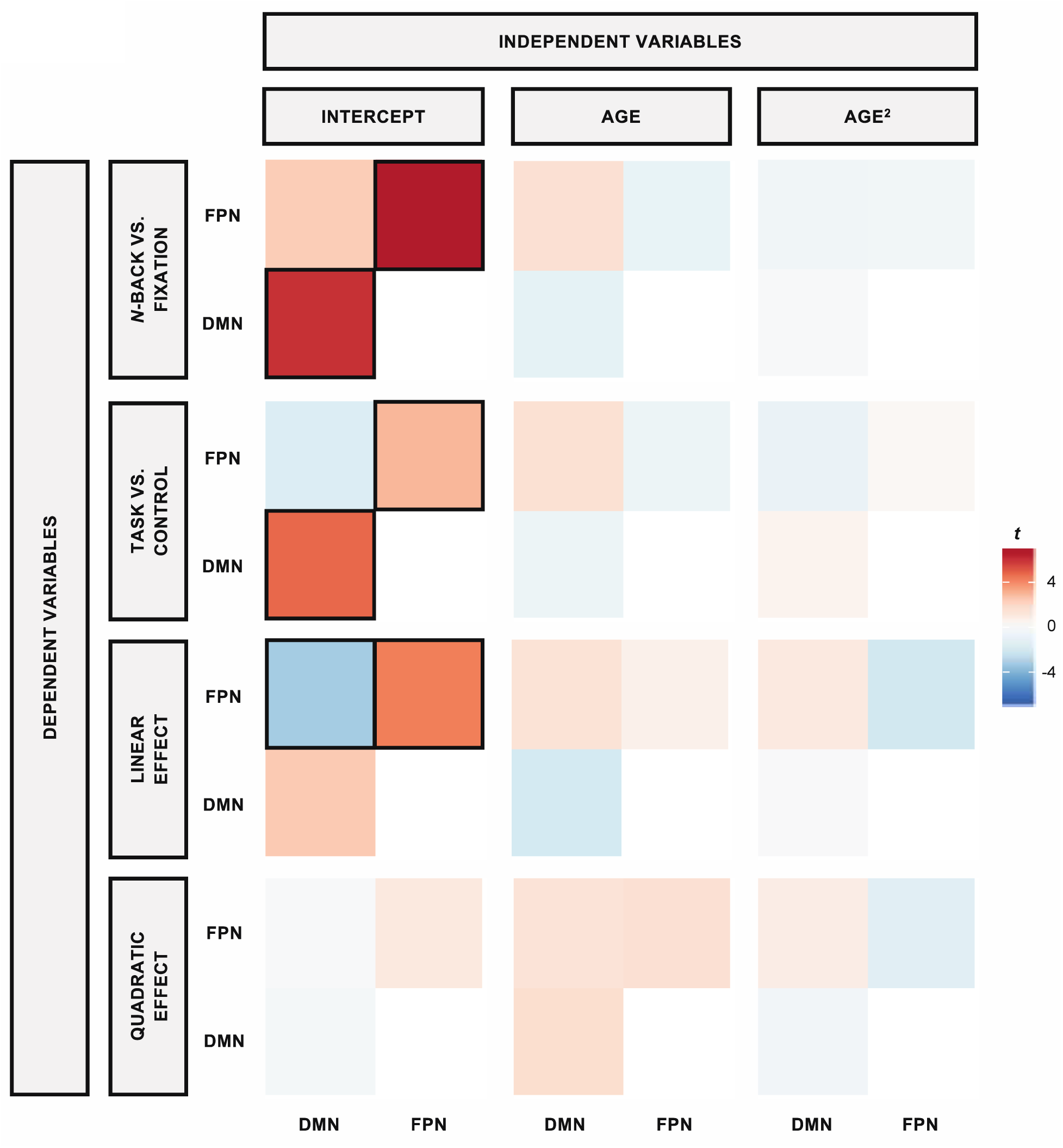
Effects of age on functional connectivity. Across all four tested models, there were no age differences in functional connectivity within- or between-fronto-parietal or default mode networks, indicating robustness of connectivity across the adult lifespan. *T*-statistics for each PPI variable for each FC network pair (DV) for linear and quadratic age are illustrated. Significant estimates, which were thresholded using an *α*_*Meff*_ of 0.004, are indicated with black boxes. Abbreviations: dependent variable (DV) and psychophysiological interaction (*PPI*).

### Effects of FC and aging on *N*-Back task performance

To examine whether FC within- and between-the FPN and DMN predicted *n*-back performance across the lifespan, *d’* was regressed on each *PPI* variable, age, quadratic age, and its interactions for each FC region. Of these analyses, only FC between the FPN and DMN during the linear effect of task significantly predicted *d’* (*b* = −0.60, *t*(164) = −4.69, *p* < .001, adjusted *r*_*p*2_ = 0.09). Specifically, *d’* increased as FC between FPN-DMN became more negatively coupled (increased anti-phase synchronization; FPN increased as DMN decreased). Furthermore, this association was moderated by quadratic age, *b* = 0.0008, *t*(164) = 2.99, *p* = 0.003, adjusted *rp*^*2*^ = 0.02. *Post-hoc* simple slopes analyses of this interaction revealed that the association between *d’* and FC between FPN-DMN during the linear effect of task was significant for individuals between the ages of 38 and 68 (*p’s* < 0.0002, adjusted *rp*^*2*^ = 0.03 to 0.09), suggesting that for middle-aged and older adults FC is a significant factor for performance, whereas it is not a significant predictor of task performance for the younger and oldest adult individuals (see Figure 5 and Table 4). Neither within-FPN nor within-DMN connectivity were associated with *d’*.

**Table 4.**
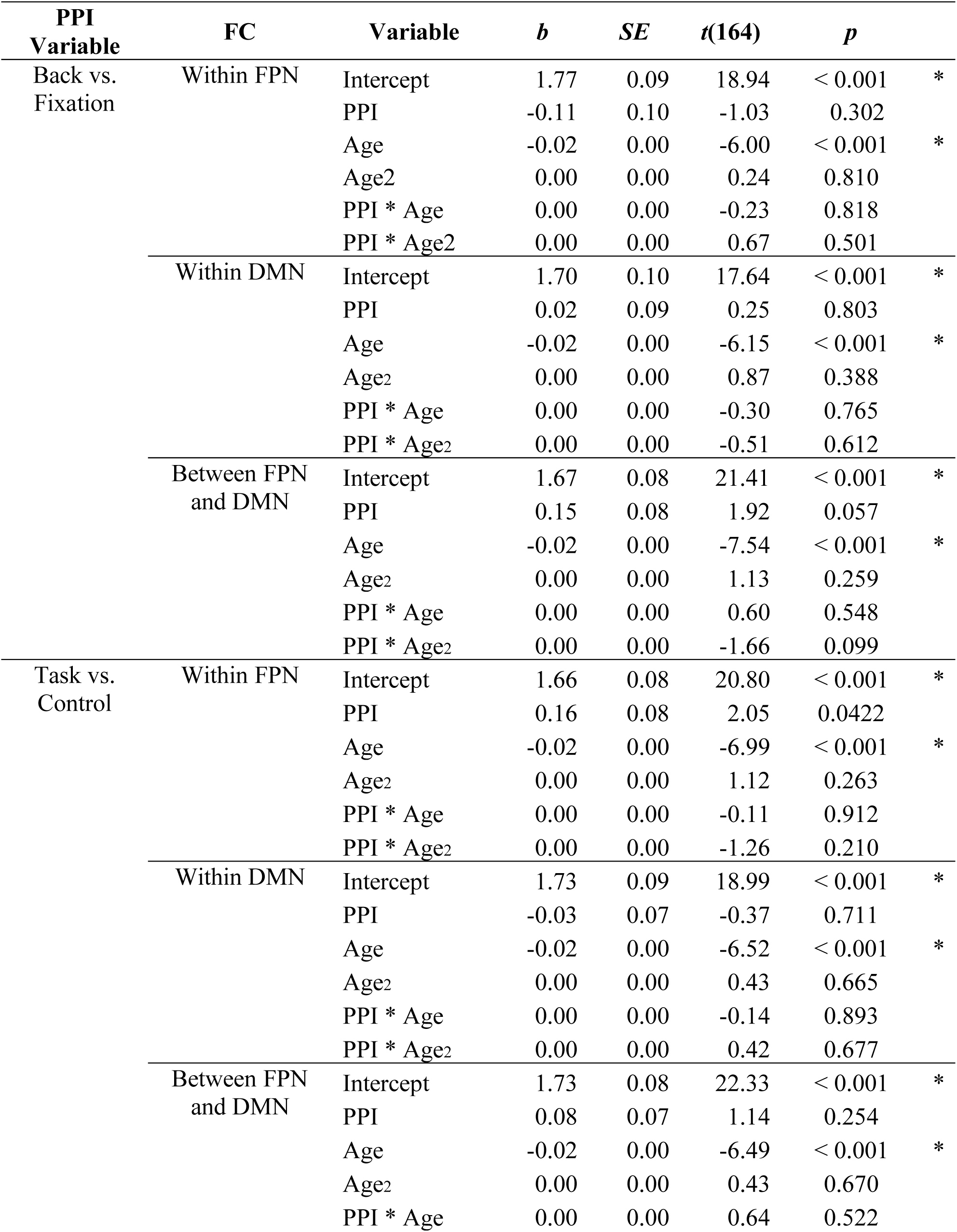
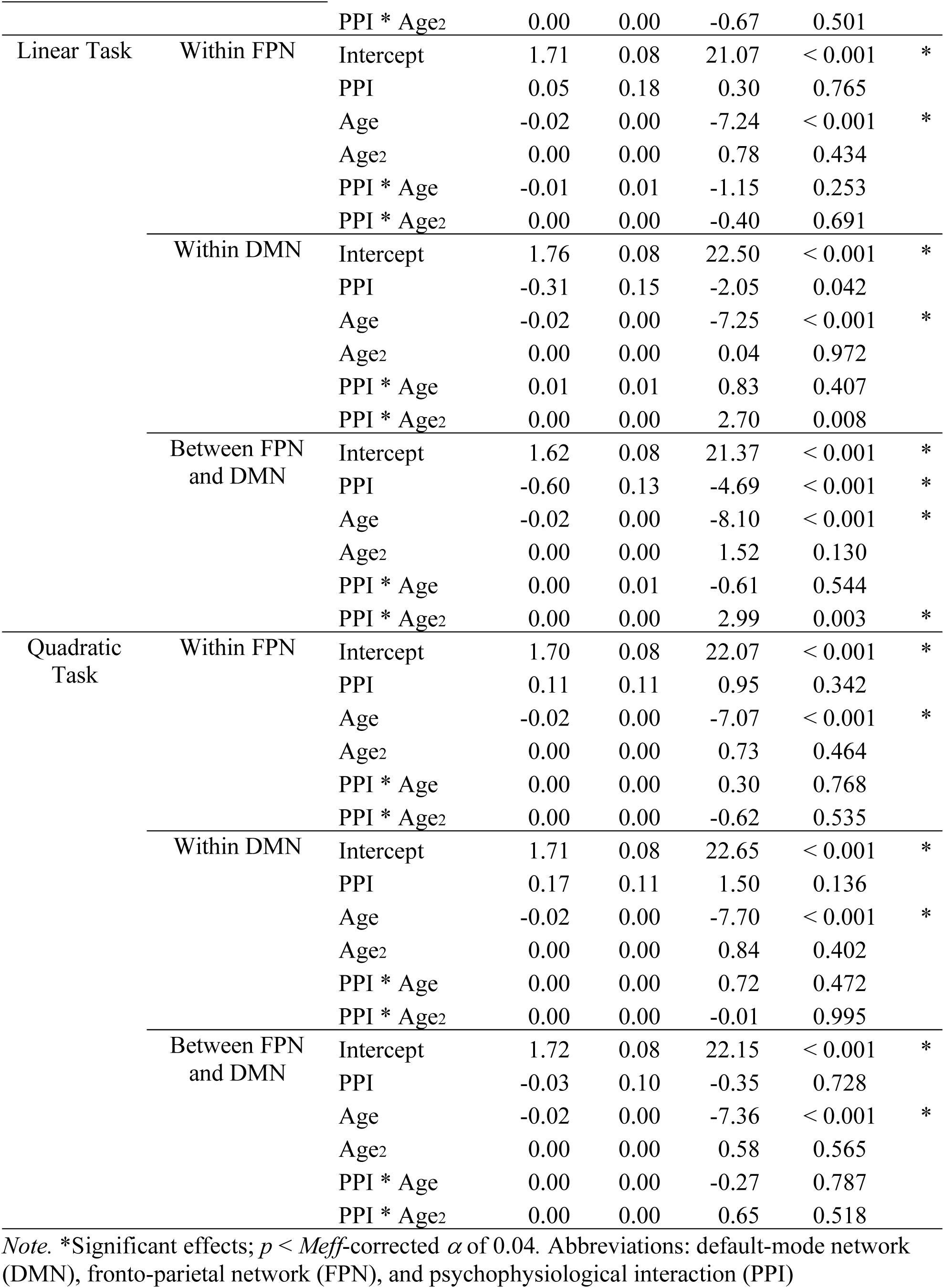
Linear and Quadratic Age, PPI, and Interactions for each PPI Variable and FC Effects on Working Memory Discrimination Index (d’)

**Figure 5.**
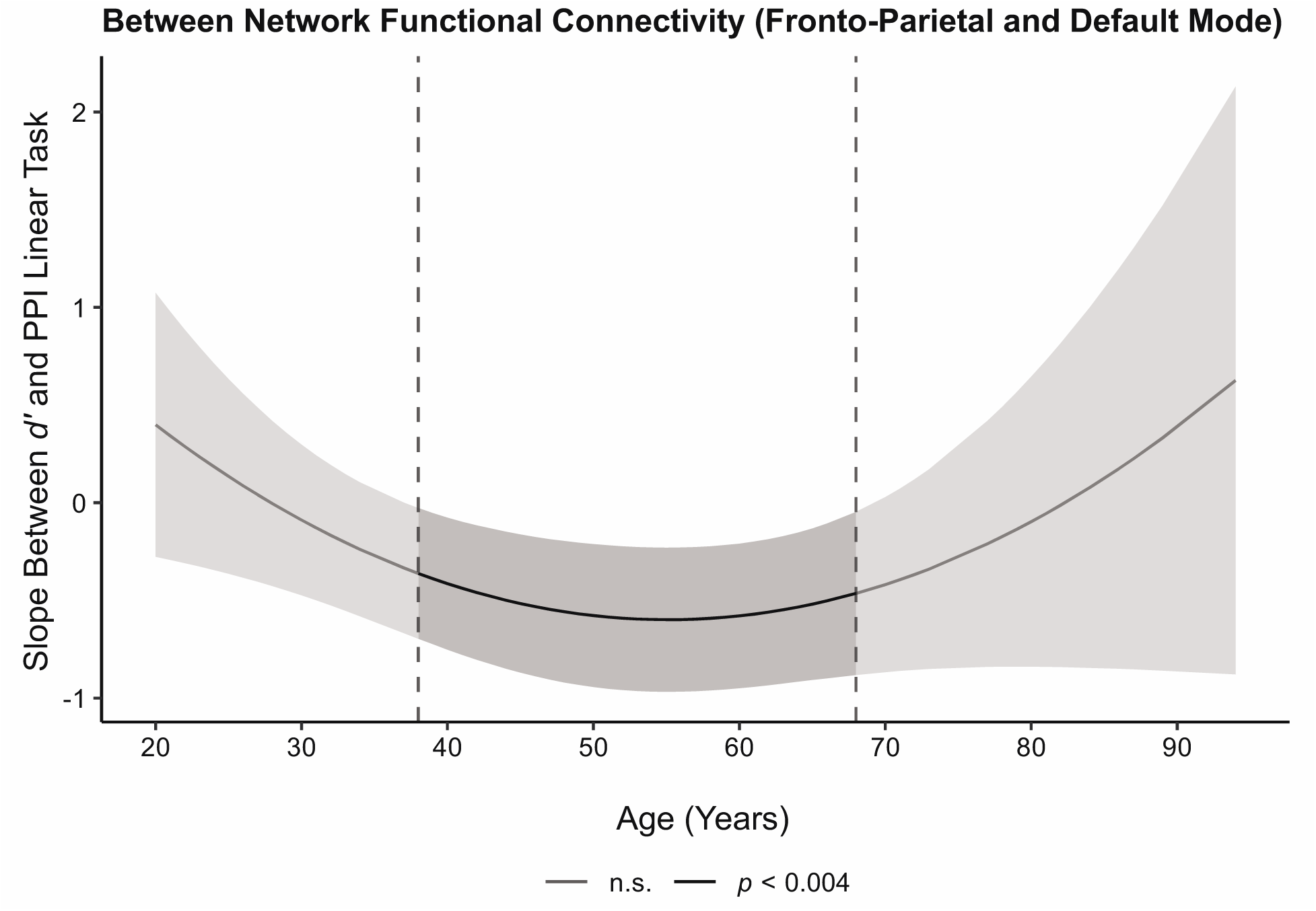
Effects of functional connectivity on cognitive performance across the adult lifespan. Working memory discriminability index (*d’*) during the in-scanner task was significantly predicted by strength of FC between FPN-DMN, but was age-dependent. *Post-hoc* analyses to decompose this significant interaction indicated stronger negative coupling between these regions (i.e., increased FPN and decreased DMN) during increasing WM load significantly predicted higher *d’* scores for middle-aged and older adults (i.e., participants between the ages of 38 and 68). Abbreviations: discriminability index (*d’*), default-mode network (DMN), fronto-parietal network (FPN), and working memory (WM).

To examine whether FC within- and between- FPN and DMN predicted out-of-scanner WM performance across the lifespan, a similar analysis was performed with digit span sequencing scores as the dependent variable. Similar results were found with only the linear effect of task significantly predicting digit span sequencing performance (*b* = −1.12, *t*(164) = - 3.32, *p* = 0.001, adjusted *rp*^*2*^ = 0.03). Specifically, digit span sequencing performance increased as FC between FPN-DMN became more negatively coupled (strengthened anti-phase synchronization). Furthermore, this association was moderated by quadratic age, *b* = 0.0027, *t*(164) = 3.405, *p* < .001, adjusted *rp*^*2*^ = 0.04. *Post-hoc* analyses revealed that the association between digit span sequencing performance and FC between FPN-DMN during the linear effect of task was significant in individuals between the ages of 42 and 58 years (*p’s* < 0.0035, adjusted *rp*^*2*^ = 0.02 to 0.04), suggesting the importance of FC for middle-aged and older adults’ WM performance, but not for the younger and oldest individuals (see Figure 6). Within-FPN and within-DMN connectivity was not associated with sequencing performance. Taken together, FC between FPN-DMN was associated with both in- and out-of-scanner WM performance, specifically, for middle and older adults, but not younger and oldest adults.

**Figure 6.**
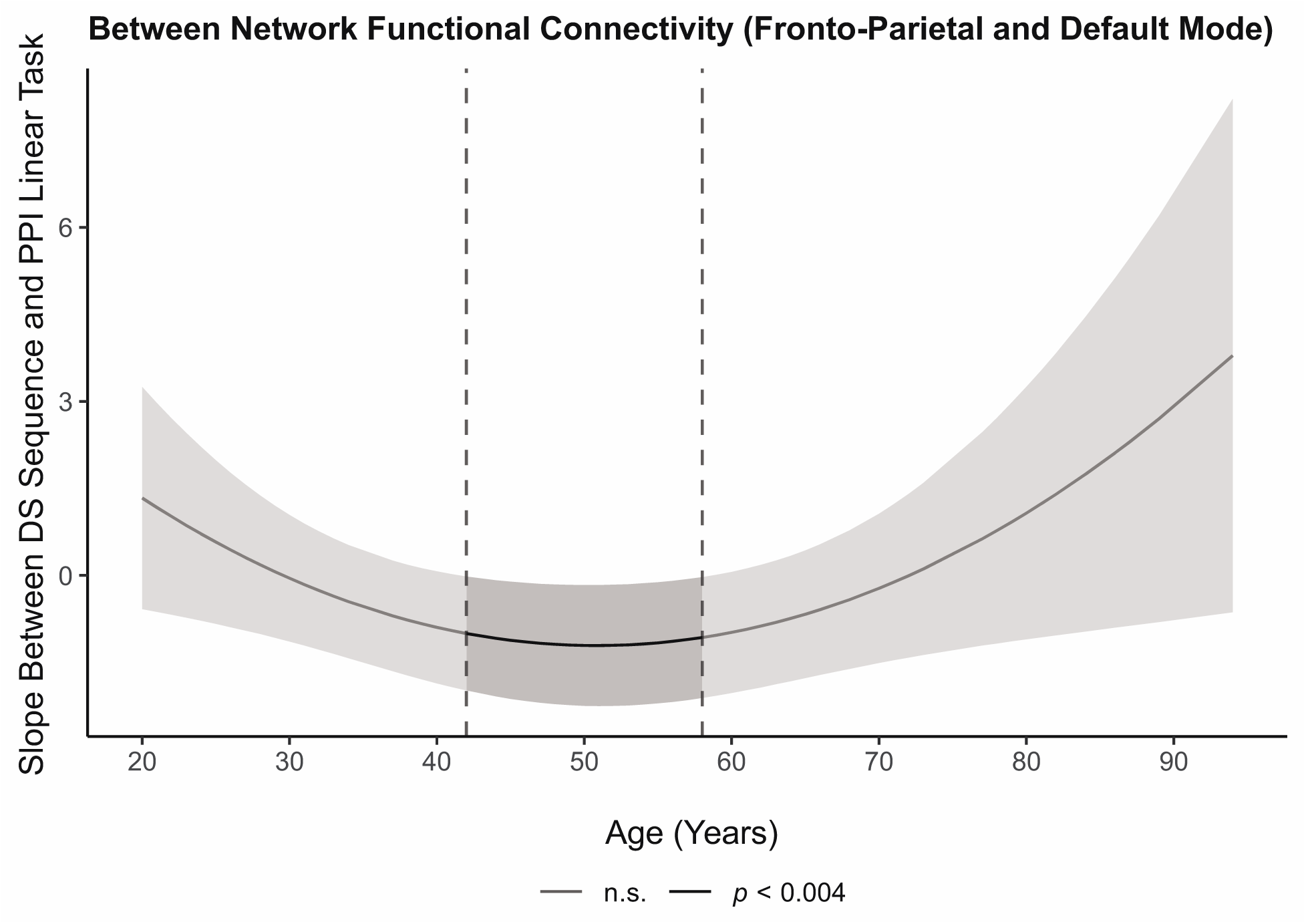
Effects of functional connectivity on out-of-scanner working memory performance across the adult lifespan. Digit span sequencing performance measured outside the scanner was significantly predicted by strength of FC between FPN-DMN, but was age-dependent. *Post-hoc* analyses revealed that stronger negative coupling between FPN-DMN was associated with better working memory performance in individuals aged 42 to 58. Abbreviations: default-mode network (DMN), digit span (DS), and frontoparietal network (FPN).

## Discussion

Here, we report that connectivity within fronto-parietal regions becomes more in-phase synchronous both when engaging in working memory on average and as WM load increases, and that connectivity within the default mode regions becomes more in-phase synchronous when engaging in WM. Furthermore, connectivity between the FP and DM networks became more anti-phase synchronous (i.e., more negatively coupled) both when engaging in WM and as load increased. Strengthening of connectivity within-FPN during WM and in response to WM load aligns with previous studies (Heinzel et al., 2014; 2017; Nagel et al., 2011; Sala-Llonch et al., 2012) that examined groups of younger vs older adults. The present findings are also in accord with a meta-analysis of PPI studies across cognitive control tasks, including working memory, compared to other cognitive domains that indicated increased FC within FP regions (i.e., dorsolateral prefrontal cortex [DLPFC] and posterior cingulate cortex [PCC]) (Smith et al., 2016). Together, these results support the general notion that regions within the FP and DM networks become functionally integrated during task engagement (Shine et al., 2016).

Interestingly, FC within- and between-FPN and DMN were robust to the effects of cross-sectional aging across the adult lifespan in the current investigation. The extant literature of age effects on FC during the *n*-back task shows inconsistent associations. Older adults have been reported to show decreased (Heinzel et al., 2014; Heinzel et al., 2017; Nagel et al., 2011) and increased FC within-FPN (Heinzel et al., 2014; Heinzel et al., 2017) compared to younger adults. Additionally, older adults have also been reported to show weakened and reversed FC between FPN-DMN (Hakun et al., 2015; Turner & Spreng, 2015) compared to younger adults. The addition of the present adult lifespan study to the inconsistent extreme age group findings in the literature suggests that there may not be true cross-sectional age effects on FC within-FPN. These working memory findings parallel work in the episodic memory domain where age effects on functional connectivity are also inconsistently reported, with some studies observing decreases in FC with aging using age group comparisons (Foster, Picklesimer, Mulligan, & Giovanello, 2016; King, de Chastelaine, & Rugg, 2018; St. Jacques, Rubin, & Cabeza, 2012; Tsukiura et al., 2011), others finding increases in aging using age group comparisons (Foster et al., 2016; King et al., 2018; Oh & Jagust, 2013; Trelle, Henson, & Simons, 2019), and some reporting age-invariance in FC (Trelle et al., 2019). Thus, for the episodic memory literature, as also demonstrated in working memory, FC differences in task sensitive regions are mixed or lack clear directionality. The lack of consistent findings in multiple literatures, as well as the absence of age effects in the current study that utilized a large adult lifespan sample, provide evidence that functional connectivity may be an age-invariant brain property.

We speculate there are several ostensible reasons for the present age-invariance compared to the mixed literature. First, there are likely additional null cross-sectional age findings during *n*-back that have not been published given the “file-drawer problem” (Rosenthal, 1979) that has persisted into a “replication crisis” (Lindsay, 2015). However, further replication, particularly in longitudinal samples that are more sensitive and more precisely capture individual aging effects are warranted (and is currently underway for the current sample). Second, our findings of age-invariance in FC may also be due to differences in sample characteristics in the current sample vs some of those in the literature. For example, in addition to the inclusion and exclusion criteria of the aforementioned studies (Heinzel et al., 2014, 2017; Nagel et al., 2011), the present study additionally explicitly excludes participants for cardiovascular disease (except for controlled essential hypertension), head trauma with loss of consciousness, and diabetes. These health conditions alone have been shown to alter task-free FC compared to healthy controls (Li et al., 2015; Yang et al., 2016; Zhang et al., 2016). Thus, we speculate that the lack of an age effect on FC may be due to our sample representing the healthy end of the normal aging continuum, whereas other samples may represent typical aging, including the influence of some common age-related comorbidities (Meusel et al., 2014). Third, the differences in age effects reported in the literature may stem from diverse methodological approaches and thresholding stringencies applied across studies. For example, the current study implemented a fairly stringent correction for multiple comparisons to mitigate Type I error, whereas most of the published aging papers utilized no alpha correction (with the exception of Heinzel et al., 2017), leaving the possibility that some effects may be spurious and influenced by Type I error. Multiple methodologies have been applied to the computation of FC including standard PPI, gPPI, granger causality, and so forth, introducing additional method variance across the literature. A well-controlled meta-analysis could help tease these effects apart.

Intriguingly, although FC may be robust to the effects of age, both positive and negative parametric BOLD modulation during *n*-back are adversely affected by aging (Kennedy et al., 2017). It is interesting to speculate what the differences in these BOLD metrics might represent. BOLD activation, and modulation of activation to difficulty reflect a magnitude-level effect of changes in oxy-deoxyhemoglobin from one state to the next, as a proxy for neurovascular coupling at the neuronal unit. In contrast, functional connectivity is a proxy for functional integration of different neuronal populations in a time-linked fashion (Friston, 2011). It is plausible that biological mechanisms induce age-related constraints or limitations on increasing magnitude of BOLD activation, whereas different mechanisms underlie functional integration, such as white matter connections across the cortex and subcortical regions. While these white matter connections are degraded with even healthy aging (Bennett & Madden, 2014; Kennedy & Raz, 2015), frank loss of axons or neuronal cell bodies is not observed in healthy aging (Liu et al., 2017). Functional integration may be maintained in this healthy sample because white matter tracts still provide an avenue for long range functional integration. In our prior reports, a significant coupled relation between positive modulation in FP regions and negative modulation in DM regions was not altered with age in this same sample of participants (Kennedy et al., 2017; Rieck et al., 2017). Thus, in separate investigations, BOLD modulation to cognitive difficulty is significantly altered with aging, however, both coupling of modulation and FC between FP and DM are age-invariant. However, changes in functional integration might be seen with further diminished white matter integrity, outside of healthy aging, as in individuals with mild cognitive impairment or Alzheimer’s disease (Wang et al., 2015; Wee et al., 2012).

In addition to characterizing how task-related connectivity behaves across the lifespan, it is also crucial to yoke this connectivity to performance (Grady, 2012). Here, we report that FC *within* FPN or DMN did not predict WM performance, rather the strengthening of FC *between* FPN and DMN as task load increased was significantly associated with better in- and out-of-scanner task performance. Middle-aged and older adults who further increased activity in FPN and simultaneously decreased activity in DMN to increasing WM load achieved higher accuracy during *n*-back performance and digit span sequencing. Regardless of FC between FP and DM regions, the youngest adults appeared to perform well, while the oldest adults performed poorer. Potentially, anti-phase synchronization (i.e., coupling) between FPN and DMN regions may not be as important to *n*-back performance in the youngest or the oldest adults compared to adults in the middle to older age range, a period theorized to be crucial for brain maintenance and/or compensation mechanisms (Cabeza et al., 2018).

This pattern of performance associated with FC between FPN-DMN regions is partially aligned with the DECHA model, which states that FPN activation coupled with DMN suppression during executive function tasks associates with better performance (Turner & Spreng, 2015). We have previously shown in this sample that greater coupling of positive and negative BOLD modulation to difficulty was associated with higher fluid intelligence (Rieck et al., 2017). Our findings also partially align with the finding that increased global functional integration during difficult external tasks such as the *n*-back paradigm is associated with effective performance (Shine et al., 2016). Additionally, previous findings from Kennedy and colleagues (2017) suggest that strengthened activation of FPN to increasing *n*-back load (positive modulation) is associated with better performance for middle-aged, older, and oldest adults but not younger adults (younger adults performed generally well regardless of strengthened activation of FP to increasing *n-*back load); while suppression (deactivation) of DMN to increasing *n*-back load (negative modulation) is associated with better performance regardless of age.

## Conclusion

In sum, the present study revealed that functional connectivity within fronto-parietal and default mode regions strengthened (became more positively correlated or in-phase synchronous) when engaging in working memory compared to control, and that FC within FP regions further strengthened as working memory load increased. Additionally, negative FC between the FP and DM was strengthened (became more anti-phase synchronous) when engaging in the WM task compared to control and as WM load increased. Notably, these patterns of connectivity were age-invariant across the adult lifespan. Importantly, however, the association of connectivity strength was linked to task performance in an age-dependent manner. Specifically, the stronger the negative coupling between FPN and DMN as task load increased, the more accurate was the task performance, selectively in middle to older-aged adults, i.e., portions of the age span that are often excluded from most “aging” samples. Further replication supporting the maintenance of functional integration across healthy cognitive and brain aging is warranted, particularly using longitudinal studies to quantify within-person change.

## Acronyms

*α*: Alpha, Type I Error Rate
*b*: Beta Estimate, Regression Coefficient
AC: Anterior Commissure
AFNI: Analysis of Functional NeuroImages
BOLD: Blood-Oxygen-Level-Depedent
CAREN: Consensual Atlas of REsting-state Network
CSF: Cerebral Spinal Fluid
*d’*: Discriminability Index
DECHA: Default-Executive Coupling Hypothesis of Aging
DCM: Dynamic Causal Modelling
DLPFC: Dorsolateral Prefrontal Cortex
DM: Default-Mode
DMN: Default-Mode Network
DS: Digit Span
EPI: Echo Planar Imaging
FC: Functional Connectivity
fMRI: Functional Magnetic Resonance Imaging
FP: Fronto-Parietal
FPN: Fronto-Parietal Network
FWHM: Full-Width Half-Maximum
GLM: General Linear Model
gPPI: Generalized PsychoPhysiological Interaction
Hem.: Hemisphere
HP: High-Pass
HRF: Hemodynamic Response Function
IPL: Inferior Parietal Lobule
ITG: Inferior Temporal Gyrus
L: Left
MCC: Middle Cingulate Cortex
*M*_*eff*_: Effective Number of Tests
MFG: Middle Frontal Gyrus
MMSE: Mini-Mental State Examination
MNI: Montreal Neurological Institute
MP-RAGE: Magnetic Prepared Rapid Gradient Echo
MRI: Magnetic Resonance Imaging
mSFG: Medial Superior Frontal Gyrus
n.s.: Not Significant
PC: Posterior Commissure
PCC: Posterior Cingulate Cortex
PPI: PsychoPhysiological Interaction
PHYS: Physiological
PSY: Psychological
R: Right
ROI: Region of Interest
*rp*^*2*^: Partial r-squared, explained variability
SMG: Supramarginal Gyrus
SPM: Statistical Parametric Mapping
*t*: *t-Statistic*
TE: Echo Time
TR: Retrieval Time
TrIFG: Inferior Frontal Triangularis
WAIS: Wechsler Adult Intelligence Scale
WM: Working Memory
z: z-Score

## Acknowledgments

We would like to thank Andy Hebrank for assistance with fMRI task programming, Asha Unni for behavioral task piloting and fMRI data collection, and Marci Horn for cognitive data collection.

## Authors’ Contribution Statement

E.E.P., K.M.K., and K.M.R. conceived and outlined the idea. E.E.P. processed, analyzed, and visualized the data and results. All authors provided critical feedback and assisted in refining the study, analysis, and manuscript.

